# Multiple adsorptions Shape Collective T-Even Phage Lysis Dynamics: Insights From an Individual-Based Model

**DOI:** 10.1101/2025.05.16.654475

**Authors:** Júlia Cabrera Cortada, Namiko Mitarai

## Abstract

Bacteria infected by T-even bacteriophages exhibit a variety of multiplicity of infection (MOI) dependent behaviours, including lysis from without (LO), cell lysis induced by an exceptionally high number of secondary adsorptions, and lysis inhibition (LIN), a phenomenon in which host cell lysis is delayed as the same bacterium adsorbs multiple phages. We propose an individual-based model that captures these MOI-dependent dynamics phenomenologically. The model takes into account the development of LO resistance (LOR) after the primary infection, and LIN is modelled by a fixed average delay in latent period per secondary adsorption with stochastic variability. The model successfully reproduces the experimentally observed multiple peaks in phage production following controlled phage addition. We extend the model to simulate batch culture growth and demonstrate that synchronized collapse of LINed culture can emerge as a collective phenomenon when LOR cells have a high, but finite, LO threshold (LOR cell lysis from withOut, LORO). Finally, we apply the model to a spatially structured bacterial colony under phage attack. We find that, if the burst size is comparable to the LO thresholds, especially LORO may play a role in colony survival due to locally elevated MOI near lysing cells. These results highlight the importance of spatial dynamics and secondary adsorption thresholds in phage-host population outcomes.

## 1. Introduction

Phages are the primary predators of bacteria, shaping microbial ecosystems and driving evolutionary arms races [1]. In marine environments, the phage-to-bacterium ratio often exceeds 1 and can reach up to 100 [2], suggesting that phages frequently face challenges in locating new hosts. This condition also increases the likelihood that multiple phages adsorb to the same bacterium. In addition, bacteria frequently grow in dense colonies or aggregates, which further elevates the probability of multiple adsorption: when a cell in such a colony lyses, the released phages are surrounded by immediate neighbors, and many of them adsorb to these nearby cells before they can diffuse to more distant targets. As a result, the neighboring cells are likely to receive multiple adsorptions [3, 4, 5]. These scenarios make it crucial to understand the consequences of Multiplicity Of Infection (MOI)-dependent interactions in phage-bacteria dynamics.

T-even phages, such as T2 and T4 that infect *Escherichia coli*, exhibit a rich repertoire of MOI-dependent behaviors (see Fig. 1 for an overview and abbreviations). When a large number of phages is adsorbed to a bacterium in rapid succession, the bacterium can undergo viron-mediated Lysis from withOut (LO)—a non-productive lysis triggered by damage to the cell envelope caused by phage tail-associated lysozymes (e.g., gp5 in T4) [6, 7, 8]. If the initial phage adsorption is not overwhelming, the host cell may establish LO Resistance (LOR) within a few minutes [6, 7]. Once in the LOR state, subsequent secondary adsorptions do not cause immediate lysis, but instead trigger Lysis INhibition (LIN) [9], extending the latent period before lysis. This extended period can last for several hours and leads to a larger phage yield upon eventual lysis [10]. After the latent period, cell will undergo the Lysis from withIN (LIN) to release phage progeny.

**Figure 1:**
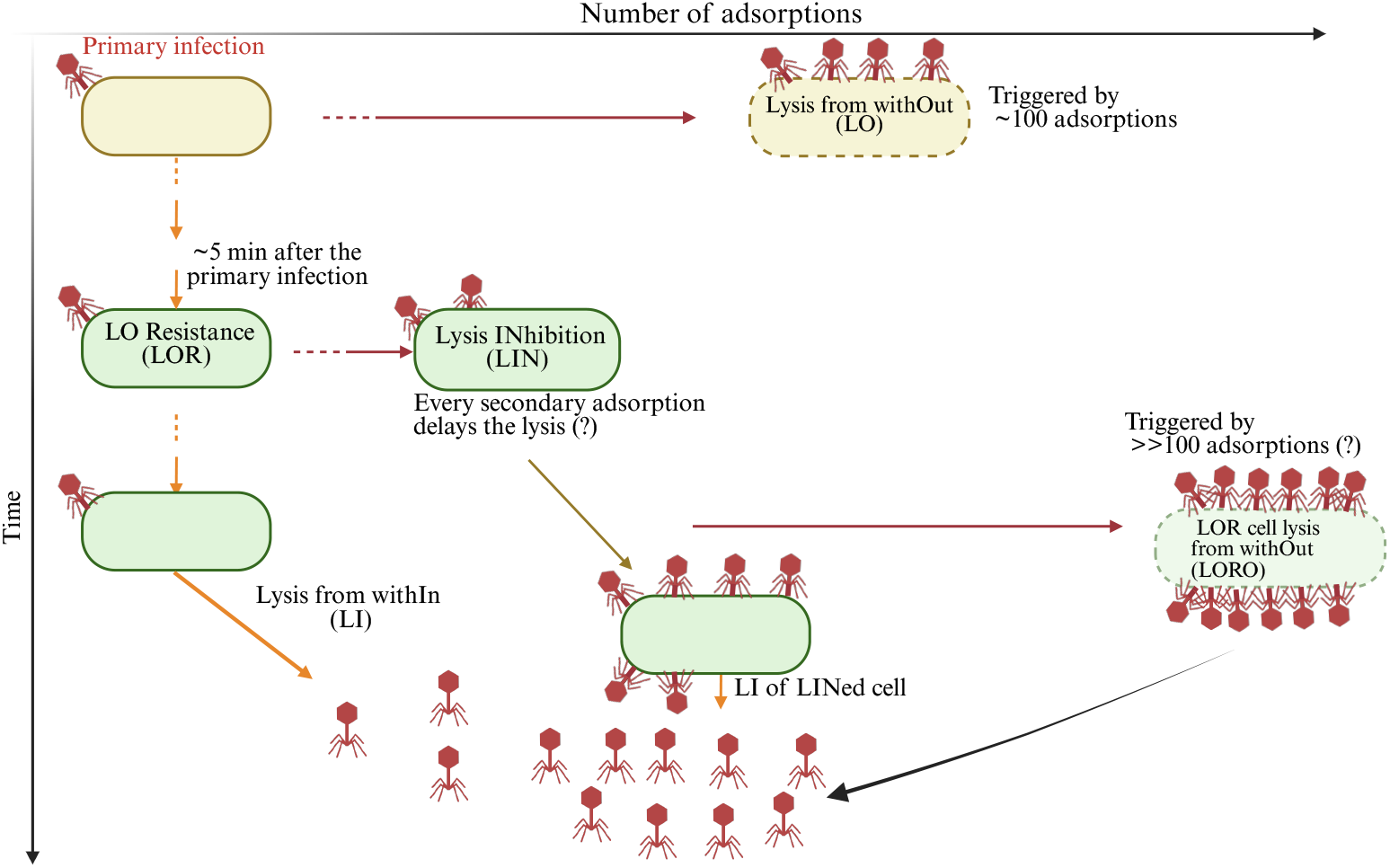
Summary of MOI-dependent behaviors in T-even phages and abbreviations. The vertical axis represents time progression, and the horizontal axis indicates the number of phage adsorptions. Yellow rounded rectangles denote infected cells before LOR is established, while green shapes represent cells that have entered the LOR state. Lysis events triggered by a high number of adsorptions (LO or LORO) are indicated by dashed outlines around the corresponding cells. Created in BioRender. Mitarai, N. (2025) https://BioRender.com/j36x932.

In liquid cultures with high concentrations of phages and infected cells, cultures undergoing LIN may suddenly clear, as a large number of cells lyse within a short time window. This synchronized burst is known as synchronized Lysis INhibition Collapse (LINC) [8, 11]. One proposed mechanism for synchronized LINC is that even LOR cells have a finite tolerance to additional adsorptions [11]—once this upper threshold is exceeded, the cell seems to undergo LO despite its previously established resistance. We refer to this as LOR cell lysis from withOut, or LORO.

While the molecular mechanisms underlying these MOI-dependent behaviors have been extensively studied [10, 7, 12, 13, 14, 15], their ecological and population-level consequences are less well understood. Among these, LIN has received the most attention, particularly with respect to its evolutionary advantages [16, 11, 17]. A high MOI suggests a scarcity of uninfected hosts, making it advantageous for a phage to delay lysis and maximize progeny output. In addition, lysis-inhibited cells continue to adsorb and neutralize additional phages, potentially reducing competition from other phage types. These hypotheses have been explored through a mathematical model of phage-bacteria population density dynamics that includes simplified versions of LIN, assuming a single secondary adsorption induces a fixed delay in lysis [18].

However, we still lack a unified theoretical framework that captures the full spectrum of MOI-dependent behaviors, including LO, LOR, LIN, and LORO, and their effects on phage-bacteria population dynamics. Experimental studies have shown that both the number of secondary adsorptions and their timing strongly influence the course of LIN. For instance, Bode [19] observed distinct peaks in phage production approximately 5 minutes apart, suggesting that each secondary adsorption adds about 5 minutes of delay and that ongoing secondary adsorption is required to maintain LIN. Capturing such dynamics requires an individual-based model that can track individual cells and their adsorption histories.

In this study, we develop an individual-based model to investigate MOI-dependent behaviors of T-even phages. In contrast to population-level models based on density variables, which represent cells with different MOIs as distinct subpopulations [20, 21, 18], individual-based models allow direct tracking of each cell’s adsorption history, including the number and timing of secondary adsorptions. Our framework incorporates stochastic adsorption and lysis timing, reproducing the complex dynamics of LIN observed in experiments [19]. We further show that introducing a threshold for LORO can explain synchronized LINC as an emergent collective phenomenon [8, 11]. Finally, we extend the model to a lattice-based spatial environment and demonstrate that local increases in MOI due to phage bursts can trigger LO and LORO in neighboring cells, which may substantially impact the survival probability of dense bacterial colonies [22].

## 2. Model

### 2.1. Individual-based model of T-even phage infection in a well-mixed environment

We model phage–bacteria interactions in a well-mixed liquid culture of volume *V*. Each bacterium is represented as an individual agent, indexed by *i* = 1, 2, · · ·, *N*, where *N* is the total number of bacteria in the system and varies over time due to cell division and lysis. The number of free phage particles, denoted by *P*, is treated as an integer-valued variable that also changes dynamically through phage adsorption and the release of progeny during lysis events. Each bacterial agent takes a discrete internal state that captures its current infection and growth status. These states encode both the cell’s progression through the phage infection cycle and its MOI- and time-dependent susceptibility to lysis from without (LO), resistance to LO (LOR), lysis from within (LI), lysis inhibition (LIN), and LOR cell lysis from without because of too high number of infection (LORO). These states and transitions are described in detail below and summarized schematically in Fig. 2.

**Figure 2:**
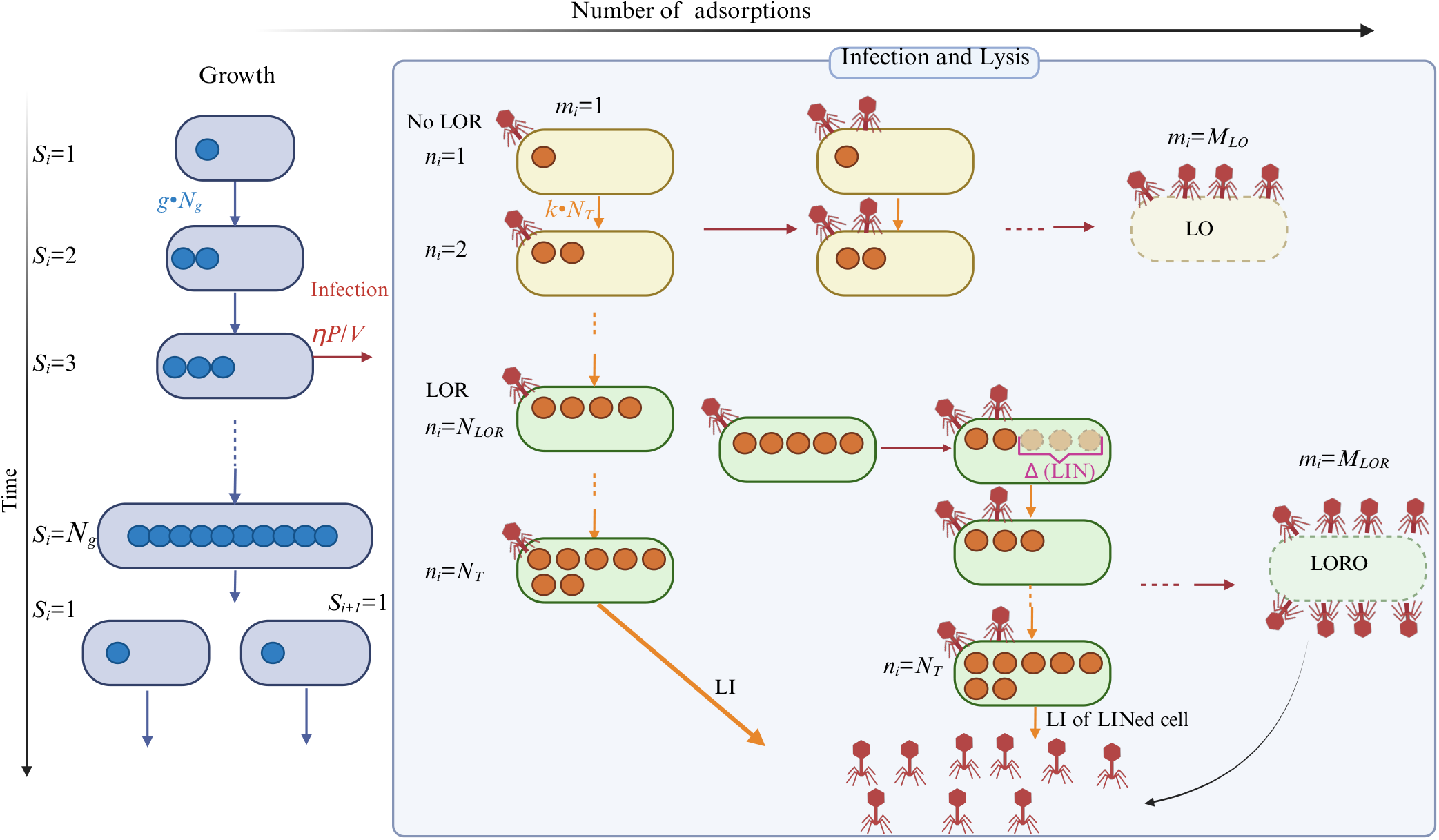
Schematic figure of the individual-based model of T-even phage and host bacteria interaction. Blue filled circles in uninfected cells indicate the growth-stage variable *S*_*i*_, which increases at a constant rate and the cell divide when *S*_*i*_ reaches *N*_*g*_. Orange filled circles in infected cells show the “timer” variable *n*_*i*_, which increases at a constant rate after infection and is reduced by Δ upon secondary adsorption once LOR is established. A cell undergoes LI when *n*_*i*_ reaches the threshold value *N*_*T*_. See main text for details. Created in BioRender. Mitarai, N. (2025) https://BioRender.com/a15g251

In the following, we introduce the rules that govern these variables’ time evolution. The summary of parameters appears in the model is summarized in Table 1.

**Table 1:** Parameters and their values.

| Parameter | Description | Default value | Source |
| --- | --- | --- | --- |
| $g$ | bacteria doubling rate | $1/30 \text{ min}^{-1}$ | typical fast growth |
| $N_g$ | growth steps<br>(model parameter, integer) | 10 | $\frac{1}{\sqrt{N_g}} \sim 30\%$ fluctuation in doubling time |
| $\eta$ | phage adsorption rate | $2 \times 10^{-9} \text{ ml/min}$ | $5 \times 10^{-10} \text{ ml/min}$ [23] to $5 \times 10^{-9} \text{ ml/min}$ [19] |
| $t_0$ | Eclipse time | 15 min | [24, 25] |
| $1/k$ | latent period length<br>(no secondary adsorption) | 27 min | [19]. cf. [23, 25] |
| $\alpha$ | phage production rate | 12.5 /min unless otherwise noted | From single infection burst size 150 [19, 25], cf. [23] |
| $\beta_{max}$ | Maximum burst size | 500 phages/cell | [24] obtained 400 to 700 phages per LINED cell |
| $M_{LO}$ | LO threshold | 100 phages | [26, 6] |
| $N_T$ | Threshold for LI<br>(model parameter, integer) | 250 | Varied to reproduce the fluctuation in [19] |
| $\frac{N_{LOR}}{k \cdot N_T}$ | LOR development time | 10 min | [6] |
| $N_{LOR}$ | (model parameter, integer) | 92 | consistency with above |
| $\frac{\Delta}{k \cdot N_T}$ | LIN duration per adsorption | 5 min | Based on [19] |
| $\Delta$ | (model parameter, integer) | 46 | consistency with above |
| $M_{LOR}$ | LO threshold for LOR cell | 150 phages | Varied to be consistent with LINC time scale [11] |

#### Growth of an uninfected cell

We assume that uninfected cells divide with an average growth rate *g*, incorporating intrinsic stochasticity in division timing. Each cell *i* is assigned a discrete growth stage variable *S*_*i*_. When a new cell is born, *S*_*i*_ = 1, and it increases incrementally at a constant rate *g* · *N*_*g*_, where *N*_*g*_ is an integer controlling the variability. The cell with *S*_*i*_ = *N*_*g*_ divides at the rate *g* · *N*_*g*_. Upon division, two daughter cells are created, each resetting its growth stage variable *S*_*i*_ to 1. This formulation ensures that the mean cell division interval is 1*/g*, with a coefficient of variation of 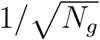.

#### Infection stage and lysis

A bacterium becomes infected by a phage at a rate *ηP/V*, where *η* is the phage adsorption rate and *P* is the current number of free phage particles in volume *V*. Each adsorption reduces *P* by one and increases the cell’s cumulative adsorption count *m*_*i*_ by one. Infected cells also track the time since the first infection, *t*_*i*_, which is used to determine the phage yield upon lysis.

We assume that a minimum time 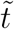 is required post-infection to produce the first viable phage progeny. After this time, the phage accumulation proceeds linearly at a rate *α*, up to a maximum burst size *β*_*max*_. An upper limit to phage production has not been established for LINed T-even phage infected cells, but we impose one for two reasons: (i) unlimited production is biologically unrealistic, and (ii) experiments with a *λ* mutant that cannot lyse from within showed that phage output saturates when chemical lysis is induced long after phage induction [27], suggesting that phage production per cell is bounded. Thus, the number of phage progeny in a cell at time *t*_*i*_is

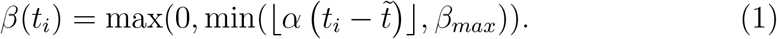

The value of *β*_*max*_ was chosen according to the existing record of T4 burst size [24]. When the cell lyses, *β*(*t*_*i*_) phages are added to the free phage pool.

We assume that infected cells no longer divide. To regulate lysis timing as a function of the secondary adsorption history, we assign each infected cell a “timer” variable *n*_*i*_, which is set to 1 at initial infection and increases incrementally unless the LIN condition is met (details given later). The rate of increase is determined by two parameters: a rate constant *k*, which sets the average latent period, and a threshold *N*_*T*_ (an integer), which determines when LIN occurs. Specifically, *n*_*i*_ increases at a rate *k* · *N*_*T*_ in the absence of further adsorptions, so that lysis occurs when *n*_*i*_ reaches *N*_*T*_, corresponding to an average latent period of 1*/k* and a coefficient of variation of 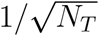. Although *n*_*i*_ plays a role reminiscent of holin accumulation in previous models of lysis timing [28], it does not represent an explicit biological entity. Unlike those models, we do not incorporate gene-expression noise; rather, *n*_*i*_ is a phenomenological construct that provides a computationally tractable way to control the variability of lysis timing.

In order to model LIN, we further implement that Secondary adsorptions reduce *n*_*i*_ to delay *n*_*i*_ reaching the threshold *N*_*T*_, but only after LOR has been established. The state transitions are handled as follows:

1. Immediately after the first infection, *n*_*i*_ is set to 1, and increases incrementally at the rate *k* · *N*_*T*_. LOR is established when *n*_*i*_ first exceeds the threshold *N*_*LOR*_. This occurs at an average time *N*_*LOR*_*/*(*k* · *N*_*T*_), which is set to be less than 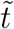, ensuring consistency with the zero-yield assumption of early LO with eq. (1). The cell is susceptible to LO until LOR is established. If *m*_*i*_, the cumulative number of adsorptions, exceeds the LO threshold *M*_*LO*_ before LOR is established, the cell is removed from the system without producing phage progeny.
2. After LOR is established, *n*_*i*_ still increases at the same rate *k* · *N*_*T*_. However, each secondary adsorption reduces *n*_*i*_ by min(*n*_*i*_−1, Δ), where Δ (an integer) is a model parameter. The minimum ensures *n*_*i*_ remains positive after the first infection. Hence, each secondary adsorption delays the time for *n*_*i*_ to reach the value *N*_*T*_.
3. After LOR, the infected cell can lyse in two ways: (i) LI: when *n*_*i*_ reaches *N*_*T*_, making the end of the latent period (with or without LIN). (ii) LORO-triggered lysis: when the cumulative adsorptions *m*_*i*_ exceed a second threshold *M*_*LOR*_ (with *M*_*LOR*_ ≥ *M*_*LO*_). In either case, the cell is removed, and *β*(*t*_*i*_) phages are released into the environment.

#### Simulation Algorithm

To efficiently simulate large populations of individuals, we employ the *tau-leaping* method [29], an approximate stochastic simulation algorithm that updates system states over a fixed time step Δ*t* based on the propensity of each event. For additional speed, we perform updates for all bacteria in parallel.

At each simulation step, we assume that there are *P* free phage particles and *N* bacterial individuals present in the system at time *t*. The following steps are performed for all bacteria **without updating** *P* during the iteration:

##### A. Phage Infection/adsorption and Lysis Dynamics

For each bacterium *i*, we sample the number of phage adsorption events *p*_*i*_ from a Poisson distribution with mean *ηP* Δ*t/V*. If *p*_*i*_ *>* 0, we execute the following:

a. Increase the cumulative adsorption count: *m*_*i*_ ← *m*_*i*_ + *p*_*i*_.
b. If this is the first infection of cell *i*, set the timer: *n*_*i*_ ← 1.
c. If the cell has not yet established LOR and *m*_*i*_ ≥ *M*_*LO*_, remove cell *i* from the system (LO event).
d. If the cell is in the LOR state:
  i. If *m*_*i*_ ≥ *M*_*LOR*_, record *p*_new,*i*_ = *β*(*t*_*i*_), remove the cell (LORO event).
  ii. Otherwise, reduce the timer variable:

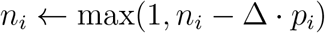

##### B. Uninfected Cell Growth and Division

If bacterium *i* is uninfected, increase its growth stage *S*_*i*_ with probability *g* · *N*_*g*_ · Δ*t*. If *S*_*i*_ = *N*_*g*_ + 1 after the update, reset *S*_*i*_ ← 1 and record a new cell division.

##### C. Timer and Lysis Progression for Infected Cells

For infected cells (*m*_*i*_ > 0), increase the timer *n*_*i*_ with probability *k* · *N*_*T*_ · Δ*t*. If *n*_*i*_ = *N*_*T*_ + 1, record *p*_new,*i*_ = *β*(*t*_*i*_) and remove the cell (LI event).

##### Final Updates

After all cells have been processed:

- Update the free phage count:

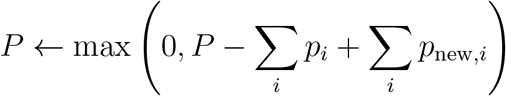
- Add the recorded number of new cells from step B to the system (each with *S*_*i*_ = 1, uninfected).
- Advance the simulation time: *t* ← *t* + Δ*t*.

This completes one simulation time step. The procedure is repeated until a final simulation time is reached.

#### Parameters

We fix most parameters that has biological meaning to the values found experimentally in phage T4 or T2 infection of *E. coli* growing in a rich medium. The values are summarized in Table 1 with units and references. Note that some of the model parameters in Table 1 do not have physical/biological meaning and are dimensionless integers.

In the result section, we vary *N*_*T*_ and Δ, which respectively control the standard deviation of the latent period and the latent period delay per secondary adsorption, to reproduce the experimental feature found in ref. [19]. We then vary *M*_*LOR*_ to be consistent with the synchronized LINC feature reported in ref. [11]. For these parameters, the finally estimated values are shown in Table 1.

The simulated volume *V* was 0.01*ml*, and the time step Δ*t* was set to be 0.01 min unless otherwise noted to ensure reasonable statistics and computational time.

### 2.2. Lattice model to simulate spatially structured environment

To investigate the impact of spatial structure, we also developed a lattice model. We are especially interested in the situation where all the surface cells of a dense colony are infected by phage, as studied in [4, 22] and illustrated in Fig. 3a. To simulate this scenario efficiently, we developed a one-dimensional model that approximates the three-dimensional system. Conceptually, this corresponds to a one-dimensional cross-section of a three-dimensional stack of bacteria, with phage predation occurring on the surface plane. This approximation captures the local structure of the colony surface while neglecting curvature effects.

**Figure 3:**
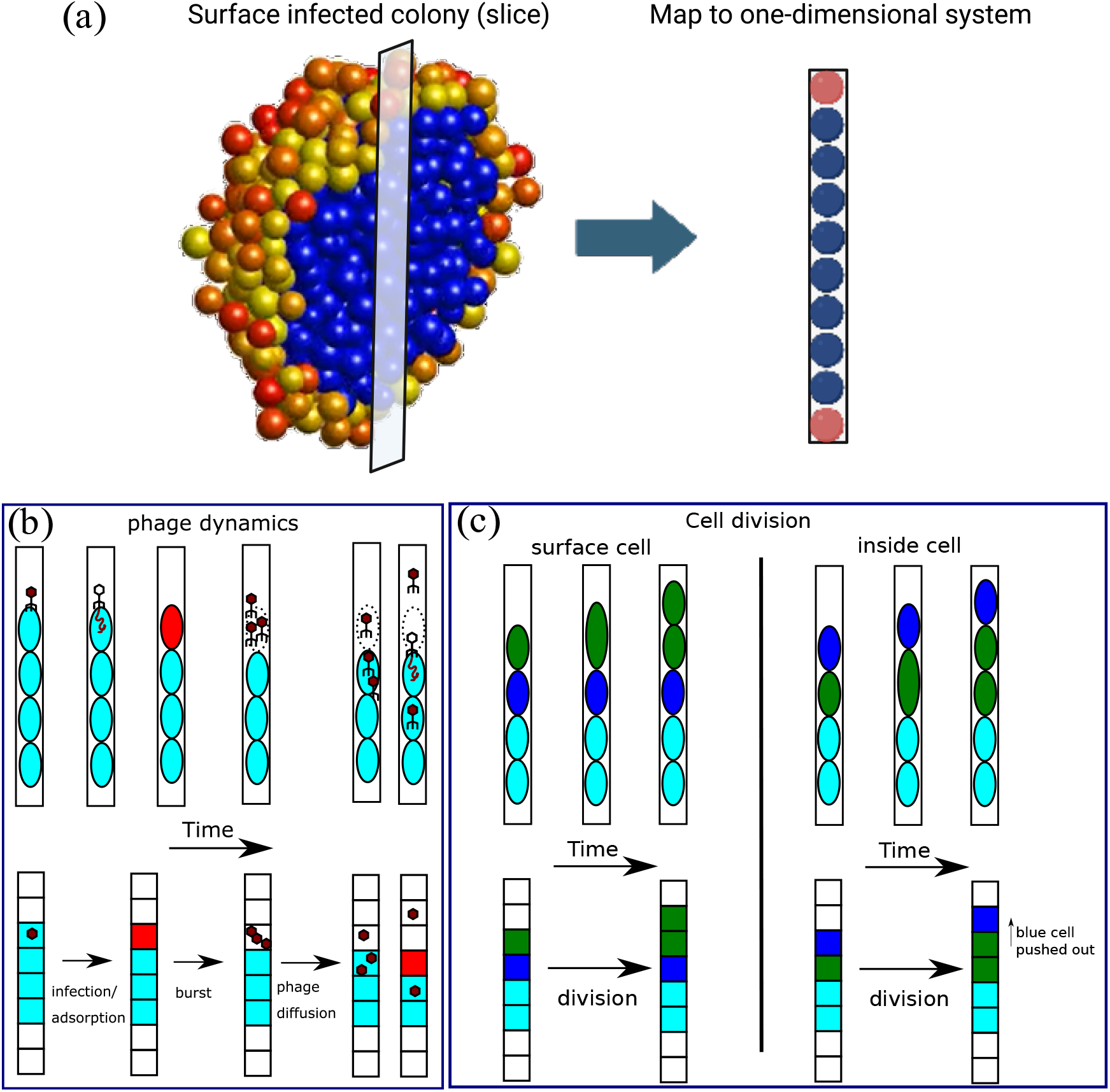
Schematic description of the lattice model. (a) Correspondence between a three-dimensional colony under phage attack and a one-dimensional lattice model. In the image, blue spheres represent uninfected cells, and yellow to red spheres represent phage-infected cells (color variation indicates time since infection). The left panel shows a slice of a spherical colony, with surface cells infected by phages. The one-dimensional lattice model (right panel) can be considered an approximation of the cutout of the cells in the colony along a line through the colony center. Created in BioRender. Mitarai, N. (2026) https://BioRender.com/o5vasg4 (b) Phage diffusion and adsorption represented. Light blue shapes represent uninfected bacteria, red shapes represent infected bacteria, and free phages are represented as brown particles. (c) Cell division and pushing are represented. Different colors are assigned to follow the dividing (green) and pushed (blue) cells.

The system consists of *L* discrete lattice sites, each of which can host at most one bacterial cell, while multiple phage particles may occupy the same site (Fig. 3b).

Phage adsorption occurs when a phage particle shares a site with a bacterium, with an adsorption rate of *η*_lattice_ = 200/min. This corresponds to a lattice site volume of *v* = 10 *µ*m^3^ when matching a bulk adsorption rate of *η* = 2 × 10^−9^ ml*/*min. Alternatively, if the site volume is closer to a single cell (*v* = 1 *µ*m^3^), the effective adsorption rate would be ten-fold lower.

Phages not undergoing adsorption move via random walks. Each phage hops to a neighboring lattice site (left or right) at a rate *k*_hop_ = 50/min. Assuming a lattice spacing of 1 *µ*m, this corresponds to an effective diffusion coefficient consistent with experimental measurements for T4 phage, approximately *D* ≈ 28 *µ*m^2^/s [30].

When interpreting the simulation as a simplification of a three-dimensional system, it is important to account for the likelihood that a phage diffuses too far to realistically return to the colony. The probability that a diffusing particle released at a distance *r* from the center of a sphere of radius *a* will hit the sphere is *a/r* [31]. Thus, phages that move beyond a distance comparable to the colony size can be considered effectively lost. To implement this simply, we track each free phage’s distance from the edge of the one-dimensional colony; if that distance exceeds the current colony size, the phage is removed from the system.

Each bacterium’s infection and adsorption dynamics, including responses to secondary adsorption and lysis, follow the same rules as in the well-mixed model described earlier.

For bacterial growth, we interpret the system as a cutout of a three-dimensional colony where nutrients are supplied from the outer surface. In such a setting, cells deeper inside the colony are expected to grow more slowly. Previous experiments in ref. [32] n three-dimensional colony growth kinetics have shown that a visible slowdown in the colony center appears only when the colony reaches about 10^6^ cells, hence a diameter of about 100 cells. Based on this, we allow uninfected bacteria located within 50 lattice sites of an empty site (typically at the colony edge) to divide at a constant rate, while cells farther inside do not divide. In a growing dense colony, this ensures that the number of actively dividing cells remains ≲ 100 (since the one-dimensional system has two edges) and that dividing cells are confined to within 50 layers from the surface.

To simulate bacterial growth so that the cells inside a colony can also divide, we assume that when an uninfected bacterium divides, all cells between the dividing bacterium and the nearest empty site are displaced one step toward that vacancy (see Fig. 3c).

Simulations are run using a fixed time step Δ*t* = 0.004 min, with a lattice size of *L* = 500 sites. At each time step, phage movement by diffusion and phage loss are updated first. Then, bacterial and infection-related dynamics at each site are updated using random sequential updates: *L* updates are performed per time step, after which the simulation time is incremented by Δ*t*. The simulation was stopped when the colony size reached 150, to ensure the colony edge was far enough from the system’s boundary to allow phages to diffuse and be lost.

### 2.3. Code availability

All simulation code was written in the Julia programming language and is available at https://github.com/namikomit/LysisInhibitionAgentBasedModel.git.

## 3. Results

### 3.1. The individual-based model with fixed average delay per secondary adsorption reproduces multiple peaks in phage production time series

We first focus on an informative experiment conducted by Bode *et al*. [19], where *E. coli* cultures at high density were infected by T4 phages at low MOI for a short time, then diluted to avoid further infections and adsorptions. After 15 minutes, the cultures were exposed to a secondary phage adsorption for 3 minutes at varying concentrations. Following this, phages were inactivated or the cultures were diluted again to prevent additional secondary adsorptions. Finally, the cultures were continuously filtered through fresh medium, and free phages in the outflow were quantified at about one-minute intervals by plaque assays.

Although the data were somewhat noisy, Bode *et al*. observed that as the secondary adsorption dose increased, phage production over time showed multiple distinct peaks, spaced roughly 5 minutes apart. This suggested that each additional secondary adsorption delayed lysis by approximately 5 minutes, though the exact delay likely depends on the secondary adsorption multiplicity.

To test whether our model can reproduce this phenomenon, we simulated the experimental setup using the following protocol: we initialized the system with 2 × 10^7^ singly infected cells per mL, simulated for 15 minutes, and then added phages at varying concentrations for 3 minutes to induce secondary adsorption. Residual free phages were then removed from the system, and from that point on, we set *η* = 0 to prevent further phage adsorption. Rather than modeling the continuous flow of fresh media explicitly, we tracked the number of phages released per minute—analogous to the experimental phage flux. Note that, in this setup, the occurrence of LO or LORO is negligible, so we are only testing the LIN-related features of the model.

We first simulated the case without secondary adsorption and compared the resulting time series (normalized to the peak height) with Bode *et al*.’s experimental data. By varying the timer parameter *N*_*T*_, which controls the stochasticity of lysis timing, we found that *N*_*T*_ ≈ 250 to *N*_*T*_ ≈ 300 yielded the best qualitative fit, (Fig. 4a). Since the experimental curve showed a longer tail and more asymmetry, we chose *N*_*T*_ = 250, which has wider distribution, as the default parameter value.

**Figure 4:**
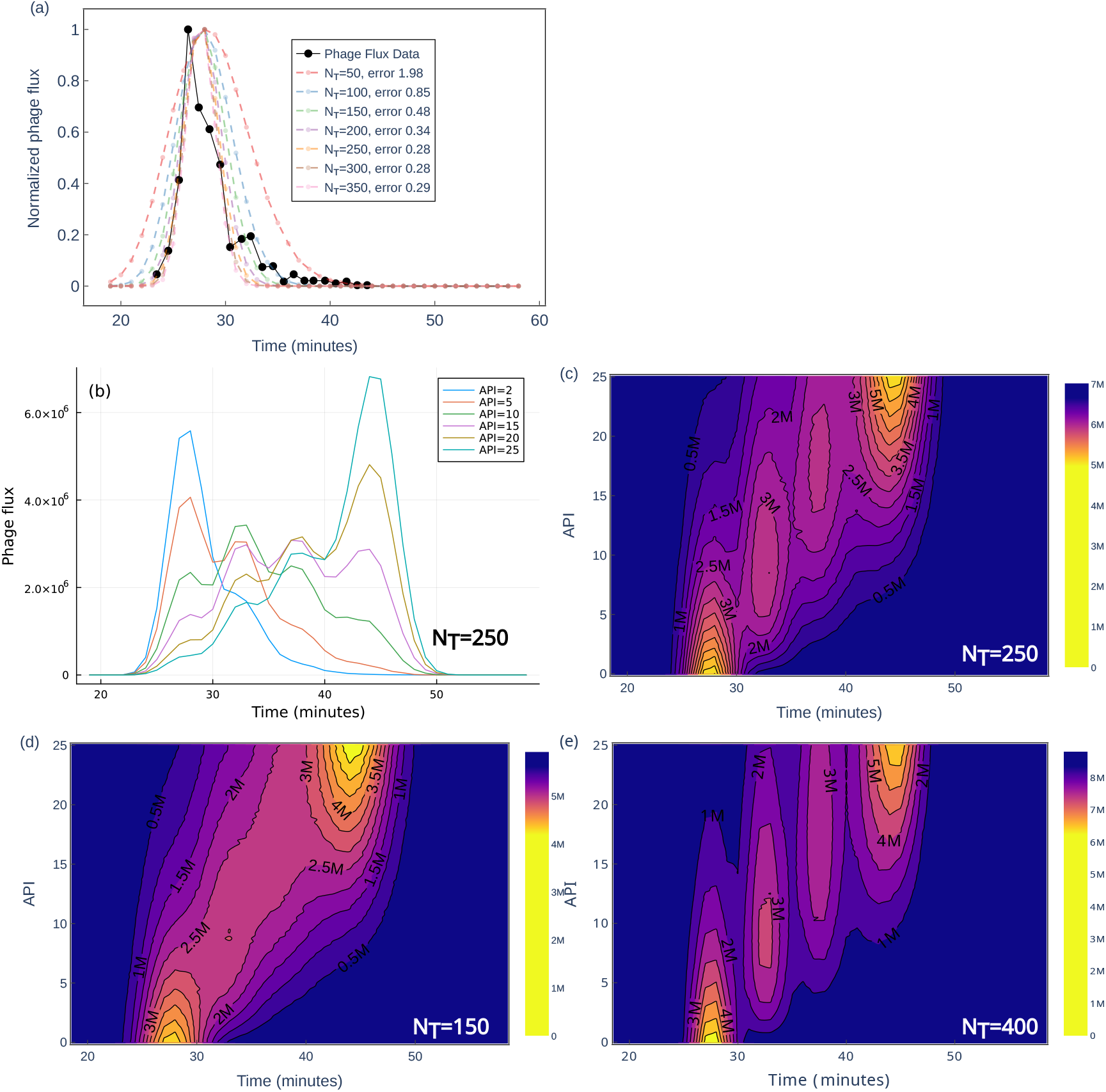
(a) Comparison of phage flux as a function of time. The experimental data was obtained from [19] Fig. 2D. The simulation was done with default parameters as described in the text, except varying the value of the number of timer molecules *N*_*T*_. All the data are normalized so that the maximum flux becomes 1. The error was quantified by the total squared distance from the data points. (b) The time series of the phage produced per min per ml at given time for the secondary adsorption *API* = 2, 5, 10, 15, 20, and 25 for *N*_*T*_ = 250. Distinct peaks are observed. (c) Heat map showing the phage produced per min per ml at given time for a given secondary adsorption *API* for *N*_*T*_ = 250. (d) The phage flux as a function of time and *API* when *N*_*T*_ = 150, which fails to have distinct peaks. (e) The phage flux as a function of time and *API* when *N*_*T*_ = 400, which shows more separated peaks than *N*_*T*_ = 250 case.

Next, we simulated the secondary adsorption protocol by varying the concentration of phages introduced during the 3-minute secondary adsorption window. Figure 4b shows the phage flux time series for selected values of the average phage input (API), defined as the number of added phages per bacterium during the secondary adsorption phase. Figure 4c presents the corresponding heat map across a continuous range of API values. For low API, the peak occurs at short times, consistent with no LIN. As API increases, multiple distinct peaks emerge, separated by roughly 5 minutes; at API = 15, four peaks can be distinguished. These peaks are progressively shifted to later times as API increases. Overall, the simulations qualitatively reproduce the experimentally observed trends. It should be noted, however, that Bode reported four peaks at API values between 1.4 and 5.7, whereas in our model a higher API was required to obtain the same number of peaks.

Finally, we tested the effect of latent period fluctuations by varying *N*_*T*_. When *N*_*T*_ is reduced to 150 (Fig. 4d), the increased stochasticity in latent period length and LIN delay blurred the peak structure, indicating that small intrinsic variability in lysis timing is crucial for observing well-separated peaks. This is further supported by the much more distinct peaks observed when increasing *N*_*T*_ to 400 (Fig. 4e).

### 3.2. Synchronized lysis inhibition collapse driven by excess secondary adsorption

Next, we simulated batch culture growth of bacteria and phage using LIN parameters determined in the previous section. For simplicity, we assume uninfected bacteria divide with an average doubling time of 1*/g* = 30 minutes and a relative standard deviation of approximately 30%. We do not model nutrient limitation, but the bacterial population does not exceed 10^9^ cells/mL in the simulations presented because of the phage predation in the setup.

Figure 5a,b shows the simulated population dynamics in batch culture. In panel (a), the vertical axis is plotted on a logarithmic scale to capture the full range of population changes, whereas in panel (b) a linear scale with a narrower range is used to highlight finer features. The dashed lines represent dynamics without a lysis threshold for LOR cells *M*_LOR_ = ∞. In this case, the system reaches a state around 50 minutes where all bacteria become infected, and the population of LINed cells decays very slowly. Meanwhile, the phage concentration fluctuates around a quasi-steady-state value.

**Figure 5:**
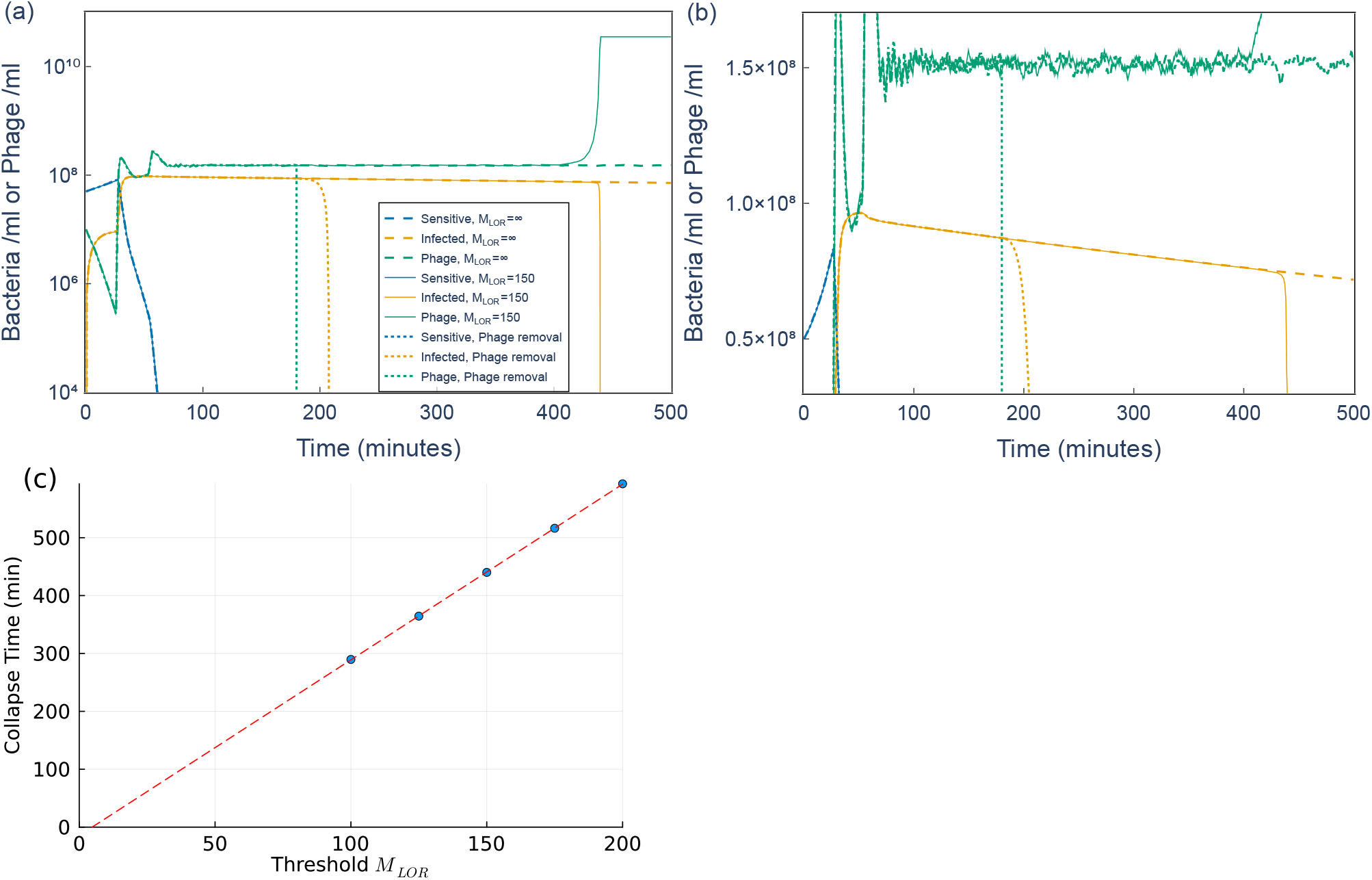
Lysis inhibition collapse simulations. (a) Population dynamics with an LO threshold for LOR cells of *M*_LOR_ = 150 (solid lines) or no LO threshold (*M*_LOR_ = ∞, dashed lines). Dotted lines indicate the case where all free phages are inactivated at 180 min, emulating the addition of an anti-phage agent to the culture [8]. Blue: sensitive bacteria (cells/ml), yellow: infected bacteria (cells/ml), green: free phages (phages/ml). The vertical axis is shown on a logarithmic scale. (b) Same as (a), but with the vertical axis on a linear scale and a narrower range. (c) Collapse timing as a function of the LO threshold for LINed cells *M*_LOR_, shown as filled circles. The dashed line indicates a linear fit with a slope of 3 min per unit of *M*_LOR_.

This quasi-steady-state phage concentration can be estimated as follows: in our model, each secondary adsorption provides a delay of about *δ* = 5 minutes in lysis. To maintain the LIN state, each bacterium must be adsorbing phage roughly once every *δ* minutes, implying an adsorption rate per cell of 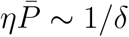, where 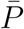 is the mean phage concentration. Using this relation, we estimate 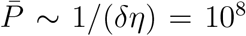 phages/mL. The actual simulation result is slightly higher, 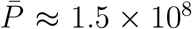, likely due to fluctuation effects neglected in the estimate.

This stable LIN state is maintained by a negative feedback loop: if phage concentration drops too low to sustain LIN, some infected cells will lyse, releasing phage and restoring the concentration. This creates mild flucuations in phage density and a slow decay of LINed cells (Figure 5b).

In reality, however, synchronized collapse of the LIN state—referred to as synchronized LINC—has been observed, with cultures clearing abruptly after a prolonged delay (e.g., ~5 hours for T4 and ~8 hours for T2) [8, 11]. Following the suggestion in those studies that LINC may result from excessive secondary adsorption causing lysis even in LOR-established cells, we simulate a finite lysis threshold for LOR by setting *M*_LOR_ = 150. The resulting dynamics (solid line, Fig. 5ab) show an abrupt increase in phage concentration around 400 minutes, followed by a sharp collapse in the bacterial population.

This synchronized collapse is driven by a positive feedback mechanism: once a cell exceeds the adsorption threshold *M*_LOR_, it lyses and releases a large number of phages due to the prolonged LIN-induced production time. These newly released phages then trigger secondary adsorption in other LINed cells, pushing them past their own *M*_LOR_ threshold. This cascade results in a collective, system-wide collapse.

Figure 5c shows how the timing of collapse depends on the value of *M*_LOR_. We observe a clear linear relationship, with collapse time increasing by roughly 3 minutes per unit increase in *M*_LOR_. This slope is comparable to the typical time between secondary adsorptions for a LIN-active cell, estimated as 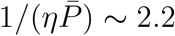 minutes.

While we lack direct experimental data to determine an appropriate value for *M*_LOR_, its effect clearly depends on other parameters such as the adsorption rate *η* and the phage burst size. For the remainder of this study, we use *M*_LOR_ = 150, as it is greater than *M*_LO_ = 100 (Table 1) and produces a biologically reasonable timescale for synchronized LIN collapse given our model parameters.

Interestingly, Abedon [8] showed that applying an anti-phage agent to inactivate free phages before collapse results in less synchronized clearing (i.e., a more gradual drop in optical density), although the timing of the collapse was similar to that in cultures without the agent. To simulate this *in silico*, we removed all free phages at 180 minutes and disabled further phage production (dotted line, Fig. 5a,b). In our model, LIN is no longer maintained, and infected cells lyse roughly 5 minutes after phage removal. The resulting population decline is more gradual than in the LINC in the *M*_LOR_ = 150 case (solid line around time 430 min), due to the lack of positive feedback to accelerate LORO. The earlier collapse in the simulation, however, suggests that additional mechanisms help maintain LIN for longer durations *in vivo*, which are not captured by the current model.

### 3.3. Role of LO, LIN, and LORO in phage attack on a bacterial colony

One natural context where secondary adsorptions frequently occur is during phage attack on a densely packed bacterial aggregate or colony. When phages infect cells in such a structure, the progeny released upon lysis are likely to be adsorbed by neighboring cells at high multiplicity [33, 3, 4, 34, 5, 22]. In particular, previous studies on a dense bacterial colony in which all surface cells were attacked by phage [4, 22] suggested that high multiplicity infection of the surface cells can protect the inner cells.

To study this scenario, we simulated a lattice model in which each site can hold at most one bacterial cell (Model section for details). We used a one-dimensional lattice, which can be interpreted as a simplified slice of a three-dimensional system, where all the surface cells of a dense bacterial colony are infected by phage (Fig. 3a), assuming symmetry in the other directions. To account for phage loss due to diffusion away from the colony [31], we tracked the distance of each free phage from the colony edge; if this distance exceeded the current colony size, the phage was removed from the system. In this way, the effective loss of phages in three dimensions is incorporated, despite the model being one-dimensional.

When the system is viewed as a slice of a three-dimensional dense colony, the growth of central cells is expected to slow once the colony becomes sufficiently large, as nutrients diffuse inward and are consumed by surface cells. Experimental study in ref. [32] has shown that dense spherical colony volumes can still grow exponentially up to 10^6^ cells (hence diameter is order of 100 cells), indicating that interior cells continue dividing at this scale. Based on this, we assume that uninfected bacteria located within 50 lattice sites of an empty site divide at a constant rate, while cells farther inside do not divide.

Phages at a given lattice site can be adsorbed by the co-localized bacterium, and their infection/adsorption and lysis responses follow the same rules as in the well-mixed model. Free phages perform a random walk on the lattice, representing diffusion, and uninfected bacteria divide by pushing adjacent cells toward the nearest empty site (see Section 2.2 for model details and parameters).

Because phages are likely to be adsorbed by surface bacteria when attacking from outside the colony, their ability to penetrate deeply into a dense colony is limited. Prior theoretical work predicts that phage infection reaches only a finite penetration depth *γ*, beyond which uninfected interior cells may continue to grow [4]. For a one-dimensional system where the bacterial doubling rate is *g*, the latent period is fixed at *τ*_*l*_, and phages penetrate to depth *γ*, the total colony size *ℓ* evolves as:

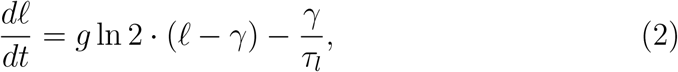

yielding a critical colony size:

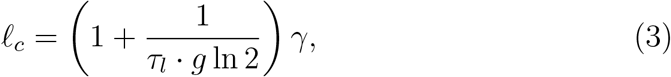

above which the colony survives and continues growing despite phage attack. However, this prediction does not incorporate the effects of LO, LIN, or LORO.

Qualitatively, LIN increases the latent period *τ*_*l*_, thereby lowering *ℓ*_*c*_ and enhancing colony resilience. In contrast, LO and LORO increase bacterial death rates, raising *ℓ*_*c*_ and diminishing survival.

To evaluate these competing effects, we simulate a scenario in which a colony of 14 cells is attacked by a single phage at each end, leaving 12 cells initially uninfected. Figure 6(a) shows a spatiotemporal plot, where uninfected, infected but not LINed, and LINed cells are visualized over time and space. Upon lysis of the initially infected cells (each releasing 150 phages), the nearest neighbors are likely to experience LO and lyse, while inner cells that have fewer phage contacts can establish LOR. These cells may subsequently enter the LIN state if further adsorptions occur, extending the latent period and allowing inner uninfected cells to divide.

**Figure 6:**
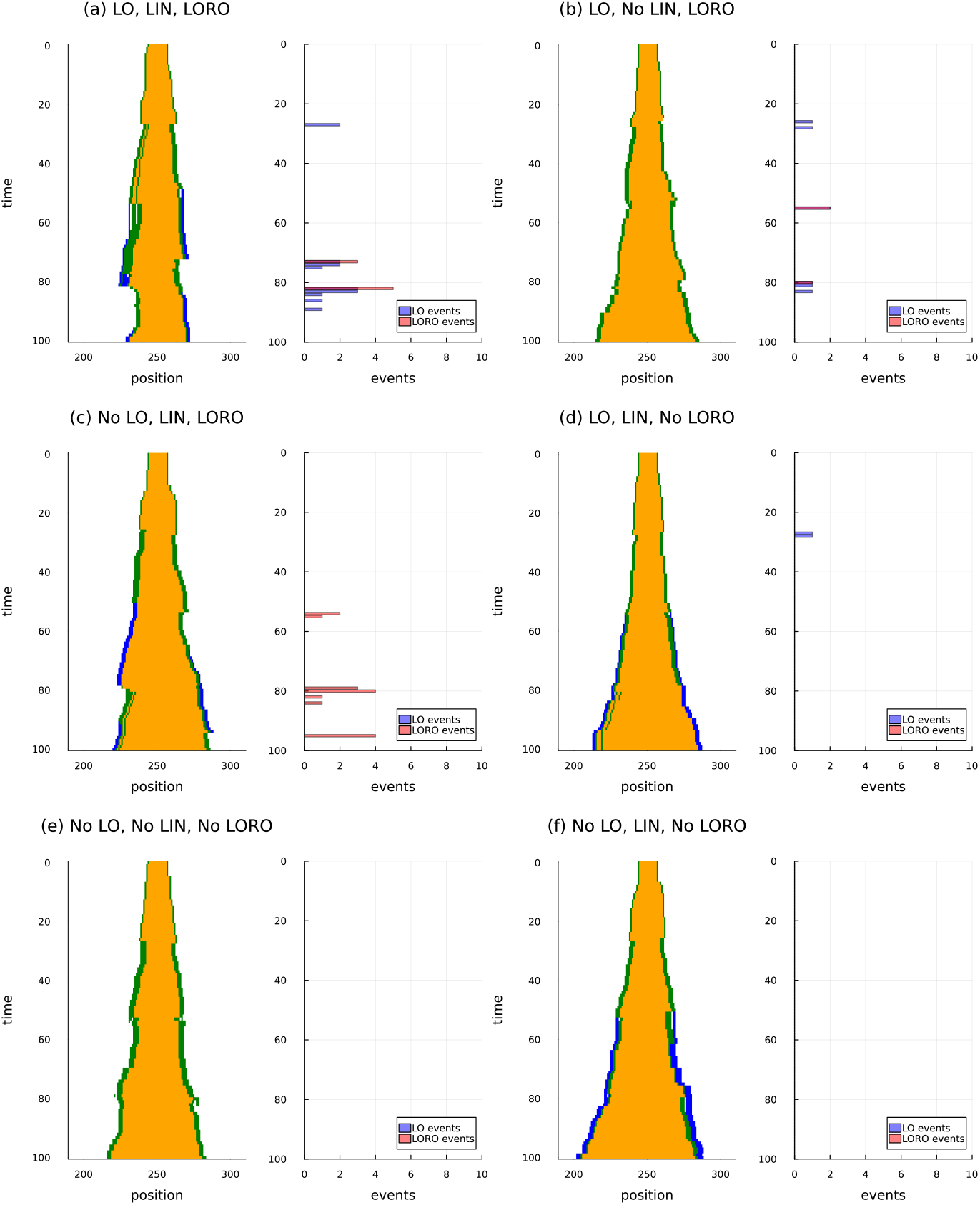
Simulation result of one-dimensional lattice model. (a-f) left: Spatiotemporal plot of the uninfected (orange), infected but not LINed (green), and LINed (blue) bacteria. Initially, 12 uninfected bacteria are placed in the middle of the system, and one infected bacteria are placed next to each end of an uninfected bacterium. The simulation is performed with a system size of *L* = 500, but only the middle 120 sites are shown. Right: The LO and LORO events per minute. (a) The full model with LIN and with LO (*M*_*LO*_ = 100) and with LORO (*M*_*LOR*_ = 150). (b) The model without LIN, with LO, and with LORO. (c) The model with LIN, without LO (*M*_*LO*_ = ∞), but with LORO (*M*_*LOR*_ = 150): In this case, the cells that has not established LOR will burst if the phage adsorption exceeds *M*_*LOR*_ and that is also counted as LORO events. (d) The model with LIN and with LO, and without LORO (*M*_*LOR*_ = ∞). (e) The model without LI, without LO, and without LORO. (f) The model with LIN, without LO, without LORO.

However, when LINed cells eventually lyse (e.g., due to insufficient secondary adsorption), their large burst size may cause nearby LINed cells to exceed the LOR threshold *M*_LOR_, triggering LORO. This creates cascades of synchronized lysis and sharp population drops, visible as large fluctuations in colony size (e.g., around time 80 in Fig. 6a). The right panel shows the number of LO and LORO events per minute, confirming that fluctuations coincide with clustered lysis events from these two mechanisms.

Figures 6(b–f) dissect the roles of each mechanism. Panel (b) removes LIN (by setting Δ = 0), reducing fluctuations but eliminating the protective effect of extended latent period. Panels (d) and (f) demonstrate that LIN contributes to protection. At the same time, (a) and (c) show that combining LIN with LORO produces large, abrupt fluctuations and suggest that the effect of LO that happens before the establishment of LOR is less important than LORO. Comparing panels (b) and (e) confirms that LO and LORO significantly suppress colony growth.

To quantify these effects, we calculate the long-term survival probability as a function of the initial colony size, assuming two infected cells are placed at the ends. Figure 7 shows that the full model (with LO, LIN, and LORO) requires substantially larger initial colony sizes to achieve survival compared to models omitting LO and LORO. In other words, while LIN alone enhances survival, its protective effects can be overridden by LO and LORO in spatially structured settings. Also, omitting LO has a smaller effect than omitting LORO alone, highlighting the importance of LORO in this setup. Notably, the model without any of these mechanisms—LO, LORO, or LIN—yields better survival for initially small colony than the full model that includes all three.

**Figure 7:**
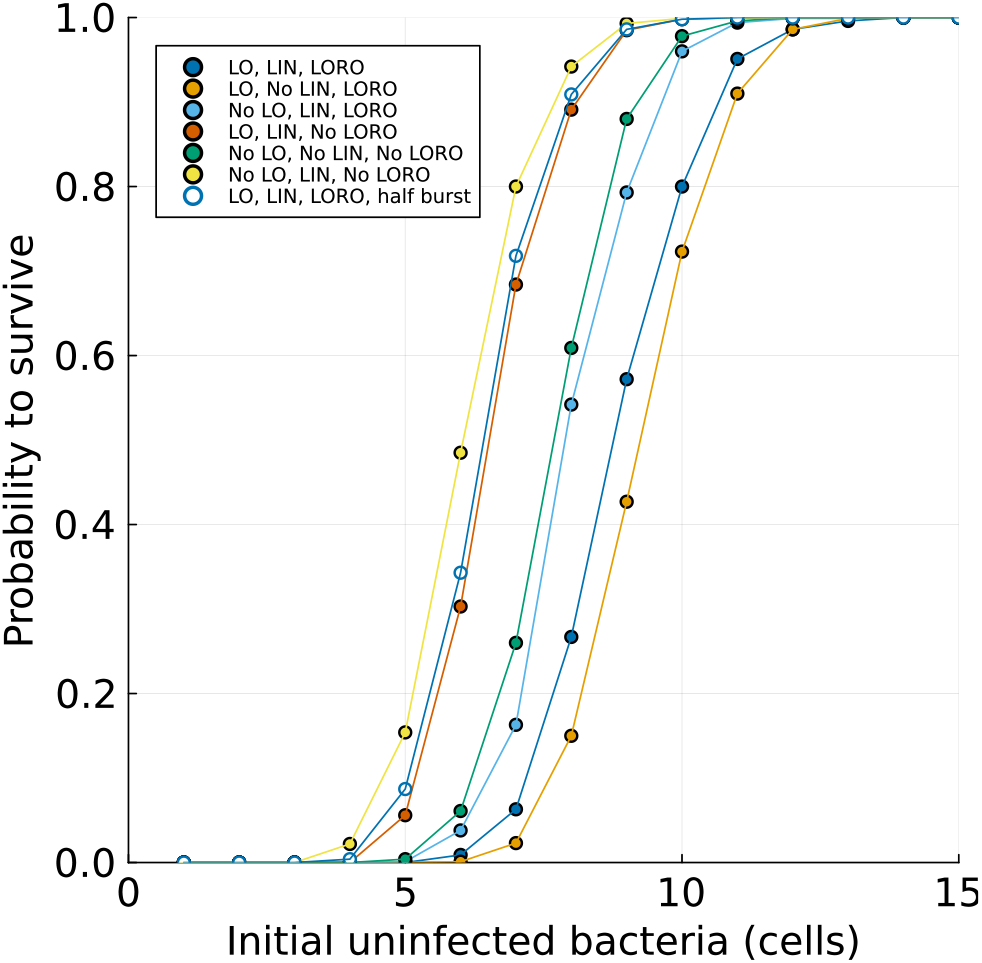
Probability for the colony to keep growing for a given number of uninfected bacteria at the start of the simulation. 1000 samples are taken for each initial condition. Survival was defined as the bacteria population reaching 150. Note that the simulations represent linear arrays of cells rather than three-dimensional colonies. Therefore, the numbers should be interpreted as a scale of colony diameter when considering correspondence to a three-dimensional case. Dark blue circle: The full model with LIN and with LO (*M*_*LO*_ = 100) and with LORO (*M*_*LOR*_ = 150). Orange circle: The model without LIN, with LO, and with LORO. Light blue circle: The model with LIN, without LO (*M*_*LO*_ = ∞), but with LORO (*M*_*LOR*_ = 150): In this case, the cells that has not established LOR will burst if the phage adsroption exceeds *M*_*LOR*_ and that is also counted as LORO events. Red circle: The model with LIN and with LO, and without LORO (*M*_*LOR*_ = ∞). Green circle: The model without LI, without LO, and without LORO. Yellow circle: The model with LIN, without LO, without LORO. Open blue circle: The full model with LIN and with LO (*M*_*LO*_ = 100) and with LORO (*M*_*LOR*_ = 150), but bust size is set to half of the default value by setting *α* = 6.25 /min.

We note that the occurrence of LO and LORO in this setup is sensitive to the relationship between the burst size and the threshold. If the burst size is substantially smaller than the threshold, phage release from a nearby infected cell may not raise the local adsorption number above the threshold. To illustrate this, we repeated the simulations with all effects included but reduced the burst size by halving the phage production rate per unit time, *α*. The resulting colony survival probability (open circles in Fig. 7) is then very close to the no-LORO case.

## 4. Discussion and conclusion

We have proposed an individual-based model for phage infection/adsorption and lysis that incorporates fluctuations in lysis timing and MOI-dependent phenomena such as lysis from without (LO), lysis inhibition (LIN), and LOR cell lysis from without (LORO). When parameters were inferred from existing literature, the model reproduced observed behaviors in well-mixed liquid culture, including multiple peaks in phage production when phages were added at controlled times, as well as synchronized lysis inhibition collapse (LINC) in LINed cultures.

Extending the model to spatially structured environments revealed that LO, especially LORO, can significantly impact colony survival, due to elevated local MOI in neighboring cells following a cell burst. These effects suggest that LO/LORO may become important to account for when assessing phage spread and bacterial survival in a spatially structured environment.

Recent experiments by Eriksen et al. [22] examined the survival of spherical E. coli colonies embedded in soft agar when challenged with high concentrations of T4 phages, revealing that a relatively large colony size was required for survival. To interpret this observation, a three-dimensional version of the model (Eq. 2) that incorporates nutrient diffusion and consumption was applied. When assuming that the phage latent period is comparable to the bacterial division time, the model estimated a phage penetration depth of approximately 27*µ*m. This is notably deeper than the ~3 *µ*m penetration depth estimate in an earlier study with phage P1vir [4], for which, to our knowledge, LO has not been reported.

Interestingly, Eriksen et al. also found an alternative parameterization for T4 infection, which assumed a latent period approximately nine times shorter and yielded a penetration depth close to 3.5 *µ*m. Although this alternative was deemed biologically implausible due to the unrealistically short latent period, our modeling results suggest that the inclusion of LO and/or LORO effects may help reconcile the observations. Specifically, these effects can accelerate effective killing by allowing a single burst event to lethally affect multiple nearby cells, thereby mimicking the impact of shorter latent periods. Thus, LO and LORO may partially explain the observed high rate of killing even under realistic lysis timing assumptions.

Our current model assumes that dead cells vanish from the system, whereas in reality, cell debris may hinder phage diffusion and adsorption, thereby reducing the efficiency of subsequent infections. Such an effect may reduce MOI and reduce the LO/LORO. Furthremore, literature suggests the existence of an upper limit in phage adsorption per bacterium, possibly due to running out of adsorption sites, though the number extends from 140 to several hundreds [35, 36, 37]. In the current modeling, we used a threshold that overlaps with this range. The determination of the upper limit can be important in understanding the role of LO/LORO in phage-bacteria interaction. This underscores the need for further refinement to better capture the complexity of T-even phage infection dynamics in dense colonies.

It is important to note that the pronounced effects of LO/LORO in spatially structured simulations depend on the assumption that the thresholds *M*_*LO*_ and *M*_*LORO*_ are comparable to or lower than the phage burst size *β*. If these thresholds were significantly higher than *β*, the frequency of LO or LORO events would be minimal, diminishing their influence, as demonstrated in the simulation with half of the default burst size.

In this analysis, we used a one-dimensional model as a simplified approximation of a three-dimensional dense colony with uniformly exposed surface cells. In three dimensions, phages released from a single burst are shared among multiple surface cells, while each surface cell can receive phages from multiple neighboring cells, effects that may partially offset each other. However, if only a subset of surface cells is infected, dimensionality may become more important. A more detailed treatment of spatial dimensionality is therefore needed to evaluate the relevance of LO/LORO in structured systems in general.

Our model also provided a few insights regarding LIN. Assuming that each secondary adsorption induces a ~5-minute delay in LIN, the multiple peaks observed in phage production experiments by Bode [19] imply that the intrinsic fluctuation in lysis timing must be substantially smaller than 5 minutes. Our model fitting suggests a coefficient of variation 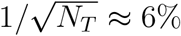, corresponding to a standard deviation of approximately 2 minutes for a 27-minute average lysis time. This low level of fluctuation is notable given the complexity of the lysis process [38].

The latent period distribution has been attracting researchers’ interest. Previous single-cell studies of latent period in temperate phage *λ* showed that fluctuations were only ~5% for a 65-minute average delay between lysogen induction and cell lysis [39]. These results were consistent with a model of holin accumulation governed by stochastic gene expression under fixed gene dosage [28]. For a virulent phage, recent single-cell observations of phage T7 infection report a much larger variation ( ~21%) for a 19-minute latent period [40], where a significant portion of the variability was due to differences in duration between adsorption and genome injection. This highlights the need for direct single-cell measurements of T4 lysis timing and LIN response to clarify how tightly T4 infection and lysis are regulated.

Our model’s prediction of synchronized LINC (Fig. 5a) features an abrupt collapse in timing, sharper than experimentally observed curves [8, 11]. One of the likely causes of this discrepancy is because our model assumes uniform parameters across all cells. In reality, variability in cell properties, such as cell size and receptor number, would lead to cell-to-cell heterogeneity in key parameters [41], including adsorption rates and the LORO threshold. Such biological noise would desynchronize cell lysis events, leading to a broader, more gradual collapse curve. Single-cell measurements of adsorption rate distributions and secondary adsorption thresholds would help refine our understanding of synchronized LINC. Moreover, the current model requires continuous secondary adsorptions to maintain the LIN state, whereas experiments show that LIN can persist even after free phages are removed at later times [8]. Together, these points highlight that the mechanisms underlying LIN maintenance remain incompletely understood, and further insight is needed to explain synchronized LINC.

Another effect that could contribute to the synchronicity of the dynamics is spatial heterogeneity, such as the formation of host aggregates. Structured populations can strongly influence phage-bacteria interactions, even when aggregates are loose and well-mixed with phages at the population level [5, 42]. This study highlights the role of LO and LORO in dense colonies, but it remains an interesting future direction to investigate how LO and LORO operate in looser structures, such as small clusters or bacterial chains, where local MOI and spatial coupling may differ.

In summary, our individual-based model captures key dynamics of phage T4 infection/adsorption, including timing fluctuations and MOI-dependent lysis phenomena such as LO, LIN, and LORO, which is difficult to take into account in concentration-based models [18]. Future experimental validation, particularly through single-cell approaches, will be crucial for deepening our understanding of phage-host interactions and refining predictive models of phage epidemics in bacterial communities.

## Acknowledgement

ChatGPT (OpenAI) was used solely for language editing and improving readability. All content and interpretations are the responsibility of the authors. This research was funded by the Novo Nordisk Foundation (NNF21OC0068775).

